# Density-dependent attributes of schooling in small pelagic fishes

**DOI:** 10.1101/2022.03.22.485264

**Authors:** Sho Furuichi, Yasuhiro Kamimura, Masahiro Suzuki, Ryuji Yukami

## Abstract

Many small pelagic fishes obligately form schools; some of these schools attain a remarkable size. Although the school is a fundamental and important ecological unit and is the site of biological interactions such as competition and predation, information on schooling processes in the field remains scarce. Here, we examined the quantitative relationships between population density and school size, the number of schools, and other school characteristics (i.e., packing density, volume, and cross-sectional area) in three species of small pelagic fishes: Japanese anchovy *Engraulis japonicus*, Japanese sardine *Sardinops melanostictus*, and chub mackerel *Scomber japonicus*. We found that school size increased almost linearly with population density, whereas the number of schools and other characteristics increased non-linearly with population density, whereby the rate of increase slowed radically as population density increased. These results indicate that, at low population densities, an increase in density causes an increase in both school size and the number of schools, whereas at higher population densities, an increase in density triggers the formation of larger schools rather than more schools. Furthermore, we found that the shapes of the relationships of all school characteristics with population density differed among species. Our results indicate that the schooling behaviour of small pelagic fishes is density-dependent, and responses to changes in density are species-specific. Our results provide insight into how biological interactions such as intra- and inter-specific competition and predator-prey interactions mediate the density-dependent processes that underlie the population dynamics and community structure of small pelagic fishes in marine ecosystems.

## Introduction

Group living is a widespread phenomenon across the animal kingdom (Krause and Ruxton 2002; Beauchamp 2014; Ward and Webster 2016), and is particularly important in fishes. Most fish species form groups at some stage of their life history, and fishes display more diverse grouping behaviour and a greater range of group sizes and shapes than other taxa (Pavlov and Kasumyan 2000; Vicsek and Zafeiris 2012).

Among fishes, the grouping behaviour of small pelagic fishes, such as sardines and anchovies, is especially distinctive and ecologically important. Many small pelagic fishes obligately aggregate to form schools, shoals, or swarms (hereafter, simply ‘schools’, although the terms are not strictly synonymous), and they sometimes form massive schools consisting of thousands or even millions of individuals (Misund 1993; Makris et al. 2006). Small pelagic fishes are also important components of marine ecosystems that link lower trophic-level plankton to upper trophic-level predators (Rice 1995; Cury et al. 2000; Bakun 2006), and the formation of schools by small pelagic fishes is likely to determine population dynamics, community structure, and energy flow in marine ecosystems through processes such as intra- and inter-specific competition and predator–prey interaction (Maury and Poggiale 2013; Maury 2017; Dalziel et al. 2021). Therefore, clarifying how and why school characteristics such as school size in small pelagic fishes vary is necessary to better understand marine ecosystem dynamics.

Population density is one of the most fundamental factors that affect schooling (Niwa 2004; Hensor et al. 2005). An increase in population density leads to a higher encounter rate between conspecific individuals or schools and hence an increased opportunity for school fusion. Therefore, mean school size is expected to increase with population density (Niwa 2004; Hensor et al. 2005). However, the formation of schools involves not only benefits (e.g., avoiding predation) but also costs (e.g., competition for food), meaning that there is likely to be an optimum school size that allows individuals to maximize their net benefit (Krause and Ruxton 2002; Ward and Webster 2016), and it has been suggested that individuals adjust their schooling behaviour toward this optimum (Sibly 1983). Therefore, school sizes might converge to a stable size or remain constant regardless of population density, rather than simply increasing in direct proportion to population density (Sibly 1983).

The relationships between population density and schooling are not simple and need to be verified by field or experimental observations. However, field or experimental studies on the relationships between these factors in small pelagic fishes remain scarce. One of the few relevant studies reported that in Atlantic herring (*Clupea harengus*), the spatial size (i.e., cross-sectional area) of schools is relatively stable, and only the number of schools increases as population density increases (Jech and Stroman 2012). On the other hand, for several small pelagic fishes in European areas, it has been reported that as population density increases, the proportion of spatially large schools increases (Petitgas et al. 2001) as does the number of schools (Brierley and Cox 2015). Therefore, in small pelagic fishes, the relationship between population density and school size remains unclear, and quantitative relationships between population density and school size and other schooling characteristics have not been well established.

The purpose of this study is to identify quantitative relationships between population density and school size, the number of schools, and other schooling characteristics in three small pelagic fishes: Japanese anchovy *Engraulis japonicus*, Japanese sardine *Sardinops melanostictus*, and chub mackerel *Scomber japonicus*. These three species inhabit the western North Pacific around Japan and are important fisheries resources that account for the majority of total fisheries catch by weight in Japan (Yatsu 2019). The stock abundance of the three species has fluctuated widely in the last decades (Yatsu 2019), which allows us to observe a wide range of population densities in the field.

First, to examine whether school characteristics (school size, number of schools, school packing density, school volume, and school cross-sectional area) increase or remain constant as population density increases and whether the shapes of the relationships are linear or non-linear (e.g., saturating), we fitted power-law models to the relationships between school characteristics and local population density at the feeding grounds of the three small pelagic fishes. Second, to examine whether the relationships between population density and school characteristics are different among species, we compared models with species-specific and species-common relationships.

## Materials and methods

### Survey

The Japan Fisheries Research and Education Agency has conducted trawl-acoustic surveys to examine the densities and distributions of small pelagic fishes every year since 2005 between September and October in the western North Pacific off northern Japan and the Kuril Islands (Fig. 1). The study area corresponds to the feeding grounds of the study species during the months surveyed. In this study, we used data from 2007, when the survey design was established, to 2021, the most recent year available. The specifications of the trawl net are as follows: opening dimensions, 30 m × 30 m; cod-end mesh aperture, 17.5 mm; tow duration, 15–60 min; ship speed, 6.5–9.3 km h^−1^ (3.5–5.0 kn). We conducted three trawl hauls per day at depths < 40 m. At each station, fish species were identified on board, and the catch was weighed by species. Up to 380 individuals of each species were randomly selected from the trawl catch and stored at −25 °C. Sampled fish were measured to the nearest 0.01 cm fork length for chub mackerel, and standard length for Japanese sardine and Japanese anchovy in the laboratory.

**Fig. 1.**
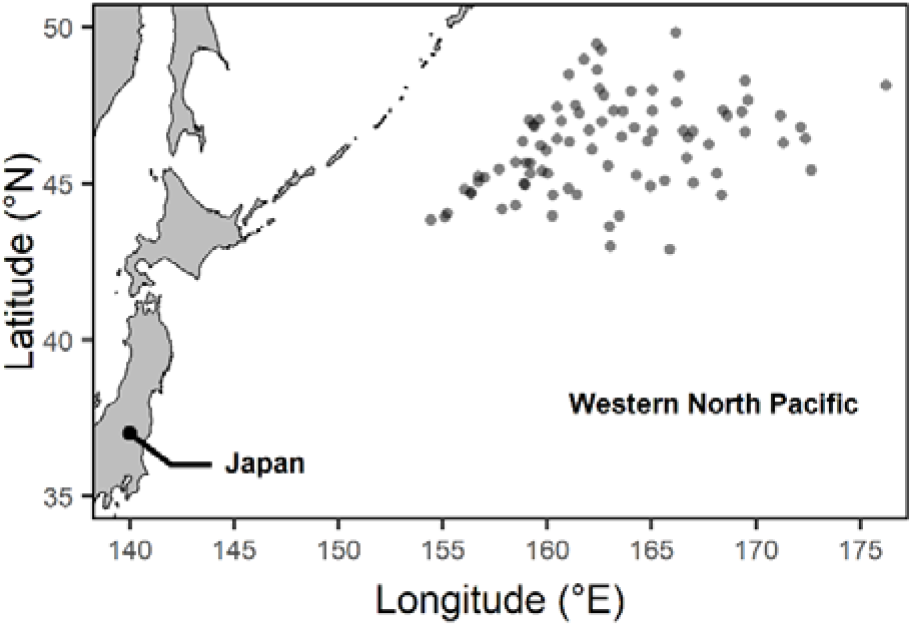
Sampling stations (translucent circles) used for trawl-acoustic surveys in the Pacific Ocean off Japan and the Kuril Islands from 2007 to 2021.

Acoustic data were collected during trawling using a Simrad EK60 scientific echosounder (SIMRAD Kongsberg Maritime AS, Horten, Norway) equipped with a hull-mounted transducer, at an operating frequency of 38 kHz. The echosounder was calibrated in accordance with standard procedures before each survey (Foote 1987). During operation, the pulse duration and ping interval were set to 1.024□ms and 0.7–1.3 s, respectively.

Conductivity–temperature–depth (CTD) water column profiles were collected at each station. The CTD profiles were used to determine sound speed (Mackenzie 1981) and absorption coefficient (Francois and Garrison 1982) for processing of acoustic data.

To analyse schools dominated by one species rather than mixed-species schools, we used only data corresponding to trawl catches where more than 80% of the trawl catch (by weight) was composed of one species (Japanese anchovy, Japanese sardine, or chub mackerel), and where catch exceeded 10 kg. Additionally, because the species of interest tend to disperse in layers at night, making it very difficult to accurately delineate separate schools (Coetzee 2000), we used only data from daytime trawls.

### Acoustic data processing

The analysis of acoustic data (volume backscattering strength [dB re 1 m^−1^]) was performed with Echoview (version 11.1; Echoview Software Pty Ltd, Hobart, Australia). To avoid potential surface noise (e.g., wave generated bubbles) and to account for near-field effects, acoustic data shallower than 7.5 m were removed. We also removed data deeper than 50 m because schools of the species of interest rarely extend deeper than 50 m in the study area (the centres of schools of the species were mainly shallower than 30 m [Table 1]). Additionally, manual inspection was used to remove any remaining undesired data (e.g., reverberation, bubble noise). To remove small spikes, two convolution kernels were applied, characterized by a 3 × 3 median filter and a 5 × 5 dilation filter (Fernandes 2009; Aronica et al. 2019). A minimum threshold of −60 dB was applied to pre-processed echograms to distinguish fishes from other particulate targets such as plankton (Jech and Michaels 2006).

**Table 1.** Characteristics of detected schools of Japanese anchovy, Japanese sardine, and chub mackerel by year. *L*_*t*_, total length; Q1, first quartile; Q3, third quartile.

| Species | Year | Total number of stations | Total number of schools | School depth (m) | | Mean $L_t$ (by station; cm) | |
| --- | --- | --- | --- | --- | --- | --- | --- |
|  |  |  |  | Median | Q1–Q3 | Median | Q1–Q3 |
| Japanese anchovy | 2007 | 1 | 5 | 24.6 | 24.1–27.5 | 12.7 | – |
| Japanese anchovy | 2008 | 3 | 21 | 22.0 | 18.0–24.7 | 11.8 | 11.1–12.2 |
| Japanese anchovy | 2009 | 12 | 53 | 26.0 | 22.4–30.9 | 12.1 | 11.6–12.4 |
| Japanese anchovy | 2010 | 10 | 68 | 20.0 | 17.8–23.9 | 12.8 | 12.5–13.2 |
| Japanese anchovy | 2011 | 13 | 92 | 21.4 | 19.2–25.4 | 10.8 | 10.4–11.8 |
| Japanese anchovy | 2012 | 8 | 34 | 19.3 | 16.7–22.0 | 13.3 | 12.9–13.7 |
| Japanese anchovy | 2013 | 2 | 21 | 26.1 | 22.6–27.6 | 13.0 | 12.9–13.1 |
| Japanese sardine | 2014 | 1 | 3 | 25.2 | 24.6–25.9 | 15.3 | – |
| Japanese sardine | 2015 | 6 | 38 | 18.9 | 16.6–22.4 | 15.3 | 15.3–15.6 |
| Japanese sardine | 2017 | 1 | 29 | 20.8 | 14.9–22.5 | 14.4 | – |
| Japanese sardine | 2018 | 4 | 61 | 16.4 | 15.7–17.6 | 13.6 | 13.4–13.7 |
| Japanese sardine | 2019 | 2 | 3 | 20.7 | 19.5–23.9 | 14.5 | 14.2–14.8 |
| Japanese sardine | 2020 | 1 | 1 | 19.4 | – | 14.5 | – |
| Japanese sardine | 2021 | 10 | 38 | 23.9 | 19.8–29.2 | 14.0 | 13.6–14.5 |
| Chub mackerel | 2010 | 1 | 3 | 14.7 | 14.4–26.0 | 21.7 | – |
| Chub mackerel | 2013 | 4 | 25 | 19.2 | 15.0–20.7 | 19.6 | 19.2–19.9 |
| Chub mackerel | 2014 | 1 | 1 | 22.3 | – | 23.1 | – |
| Chub mackerel | 2015 | 1 | 1 | 19.8 | – | 19.4 | – |
| Chub mackerel | 2016 | 1 | 4 | 21.4 | 18.0–23.0 | 18.7 | – |
| Chub mackerel | 2018 | 2 | 2 | 18.4 | 14.9–21.9 | 24.8 | 20.6–29.0 |
| Chub mackerel | 2019 | 1 | 1 | 18.4 | – | 25.8 | – |
| Chub mackerel | 2020 | 3 | 14 | 25.6 | 21.9–32.1 | 19.2 | 18.6–19.7 |

For subsequent analyses, we calculated the mean target strength (dB re 1 m^2^) of each school and at each station. Target strength indicates the acoustic reflectivity of the ensonified target (e.g., a fish). The mean target strength was calculated from the mean total length of catch samples using the following equations. For chub mackerel (Simmonds and MacLennan 2008), the equation is:

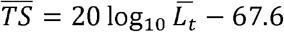

and for Japanese sardine and Japanese anchovy (Zhao et al. 2008), the equation is:

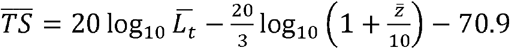

where 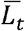 is the mean total length (cm) of a catch sample at a given station. For the mean target strength at a station, 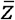 is defined as the mean depth (m) of the distribution of Japanese sardine or Japanese anchovy at the station, and for the mean target strength in a school, 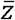 is defined as the mean depth of the school. The mean depth is the average of all sample depths at the station or in the school weighted by volume-backscattering coefficient values. Because body length was measured as fork length (*L*_*f*_) for chub mackerel, and standard length (*L*_*s*_) for Japanese sardine and Japanese anchovy, measurements were converted to total length (*L*_*t*_) by using the following formulae (Furuichi et al. 2021): *L*_*t*_ = 1.07*L*_*f*_−0.136 for chub mackerel; *L*_*t*_ = 1.13 *L*_*s*_ + 0.371 for Japanese sardine; and *L*_*t*_ = 1.12 *L*_*s*_ + 0.358 for Japanese anchovy.

We estimated local population density (ind. m^−3^) at each station by using the following formula: 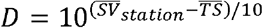, where *D* is the local population density at a station, 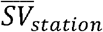 is the mean volume-backscattering strength of the water column (7.5–50.0 m) during trawling at the station, and 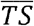 is the mean target strength at the station (Simmonds and MacLennan 2008).

Fish schools were detected and isolated from the pre-processed echograms by using the Shoals Analysis and Patch Estimation System (SHAPES) algorithm, implemented in Echoview (Barange 1994; Coetzee 2000). In the SHAPES algorithm, schools are detected by first identifying candidates, which are groups of adjacent pixels that meet minimum criteria for length and thickness. These individual school candidates are then linked together to form a larger school candidate if the horizontal and vertical distances between them are less than the specified maximum linking distances. After all linking has been carried out, schools are recognized if the final school candidates are larger than the defined minimum total school length and thickness. The detection parameters we used were as follows: minimum total school length, 4 m; minimum school height, 2 m; minimum candidate length, 2 m; minimum candidate height, 1 m; maximum vertical linking distance, 5 m; and maximum horizontal linking distance, 15 m. This set of parameters was selected based on previous studies and the resolution of the acoustic data (Coetzee 2000; Fernandes 2009; D’Elia et al. 2014).

All detected schools were classified as Japanese anchovy, Japanese sardine, or chub mackerel based on the trawl catch composition. We extracted school packing density (number of individuals m^−3^), school volume (m^3^), and cross-sectional area (m^2^) for each school. The packing density of a school was calculated as 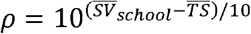, where *ρ* is the packing density of a school, 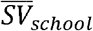 is the mean volume-backscattering strength of the school, and 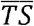 is the mean target strength in the school (Simmonds and MacLennan 2008). The volume of a school was calculated assuming a cylindrical shape as *V* = ∑_*i*_ π · (*T*_*i*_/2)^2^ · *L*_*i*_, where *V* is volume, *T*_*i*_ is the thickness of the school at ping *i*, and *L*_*i*_ is the distance from ping *i* to ping *i +* 1. School size (number of individuals) was calculated by multiplying the packing density by school volume. In addition, we also calculated the number of schools at each station.

### Statistical analysis

We examined relationships between local population density and school characteristics (school size, number of schools, packing density, school volume, cross-sectional area). To consider the possible nonlinearity of the relationships, we applied power-law models:

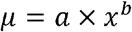

where *μ* is the mean value of a school characteristic (school size, number of schools per km, packing density, school volume, or cross-sectional area), *x* is local population density, *a* is a coefficient (>0), and *b* is a power exponent. The power exponent *b* determines the direction (positive, constant, or negative) and shape (linear or nonlinear) of the relationship. A negative power exponent (*b* < 0) means a negative relationship between local population density and a school characteristic, an exponent of zero (*b =* 0) means that a school characteristic is constant regardless of local population density, and a positive exponent (*b > 0*) means a positive relationship. Additionally, the power exponent also determines the shape of the relationship when the exponent is greater than zero: an exponent greater than one (1 < *b*) generates a convex relationship (accelerating increase with increasing local population density), an exponent equal to one (*b* = 1) generates a linear relationship (proportionality), and an exponent lower than one (0 < *b <* 1) generates a concave relationship (slowing increase with increasing local population density). The coefficient *a* serves as a simple scaling factor, meaning that the shape of the relationship will not change depending on the coefficient, but its placement will move up or down as the coefficient increases or decreases, respectively.

For school size, packing density, school volume, and cross-sectional area, we assumed that observed values follow a Gamma distribution because of their continuous, non-negative, and right-skewed distributions. Therefore, the models were defined as follows:

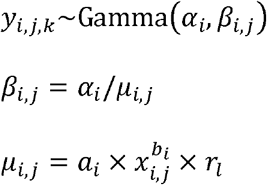

where *y*_*i,j*_ is the observed value of a school characteristic of species *i* at station *j* in school *k, *α*i* is a shape parameter (*>*0), *β*_*i,j*_ is a rate parameter (*>*0), *μ*_*i,j*_ is the expected (mean) value of the school characteristic, *x*_*i,j*_ is local population density, *a*_*i*_ is a coefficient, *b*_*i*_ is a power exponent, and *r*_*l*_ is a random effect of date *l*. Factors other than population density have also been reported to affect schooling behaviour (Borner et al. 2015; Romenskyy et al. 2020; Zheng and Fu 2021), and these factors may cause spatiotemporal autocorrelation in school characteristics. To consider this spatiotemporal autocorrelation, survey date was incorporated into the model as a multiplicative random effect following a normal distribution with mean 1 and standard deviation *σ*_*r*_.

For the number of schools, we assumed that observed values followed a negative binomial distribution because the data were count data characterized by a large variance. Therefore, the model was defined as follows:

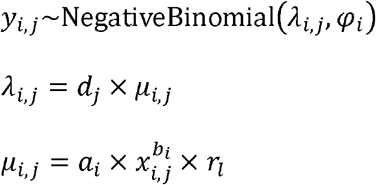

where *y*_*i,j*_ is the observed number of schools of species *i* at station *j, λ*_*i,j*_ is a location parameter (*>*0), *ψ*_*i*_ is a dispersion parameter (*>*0), *d*_*j*_ is sampling distance (km), *μ*_*i,j*_ is the expected (mean) number of schools per km, *x*_*i,j*_ is local population density, *a*_*i*_ is a coefficient, *b*_*i*_ is a power exponent, and *r*_*l*_ is a random effect of date *l*. To consider any spatiotemporal autocorrelation, survey date was incorporated into the model as a multiplicative random effect following a normal distribution with mean 1 and standard deviation *σ*_*r*_.

Furthermore, we investigated whether the relationships between local population density and school characteristics are different or common among species. In addition to the above models assuming species-specific relationships, we also constructed models assuming species-common relationships, where the coefficient and power exponent were same among species, that is, *a*_*anchovy*_ = *a*_*sardine*_ = *a*_*mackerel*_ and *b*_*anchovy*_ = *b*_*sardine*_ = *b*_*mackerel*_. To compare models, the widely applicable Bayesian information criterion (WBIC) was calculated for each model (Watanabe 2013).

We fitted the models in a Bayesian framework using Markov chain Monte Carlo methods via No-U-Turn sampling with the software STAN, which was called from R (R Development Core Team 2020) using the package ‘rstan’ (Stan Development Team 2020), to obtain posterior parameter distributions. We used weak informative priors (Cauchy or half-Cauchy distribution with location parameter 0 and scale parameter 5) for all parameters except *σ*_*r*_. For *σ*_*r*_, a normal distribution with mean 0 and standard deviation 0.2 was used as the prior distribution because of the difficulty in converging. We used four chains, each of which had 40,000 iterations including a burn-in of 10,000 iterations with a thinning interval of 10, resulting in 12,000 values for the posterior distribution of each parameter. Convergence of Markov chain Monte Carlo algorithms was checked by using the potential scale reduction statistic 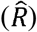 and effective sample sizes (Carpenter et al. 2017); for all parameters, the 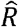 was less than 1.01 and effective sample sizes were greater than 400.

## Results

In total, data from 88 stations and 518 schools were obtained (Table 1). A wide range of local population density (ind. m^−3^) was observed ranging from 1.18 × 10^−4^ to 0.41 for Japanese anchovy, from 1.26 × 10^−4^ to 0.58 for Japanese sardine, and from 4.6 × 10^−6^ to 0.016 for chub mackerel.

### School size

For all species, school size tended to increase as local population density increased, and the power exponents were greater than zero, indicating positive relationships (Figs 2a and 3, and Table S1). In each case, mean school size increased almost linearly with local population density (Fig. 3). The estimated power exponents were relatively close to one (although slightly lower in chub mackerel than other species), but the credible intervals did not include one (Fig. 2a and Table S1). This indicates that school size increased almost linearly with local population density, but the rate of increase slowed slightly as local population size increased. The model with species-specific relationships between school size and population density had a lower WBIC than the model with species-common relationships (species-specific, 37.8; species-common, 60.6), indicating that the relationships differed among species (Fig. S1a).

**Fig. 2.**
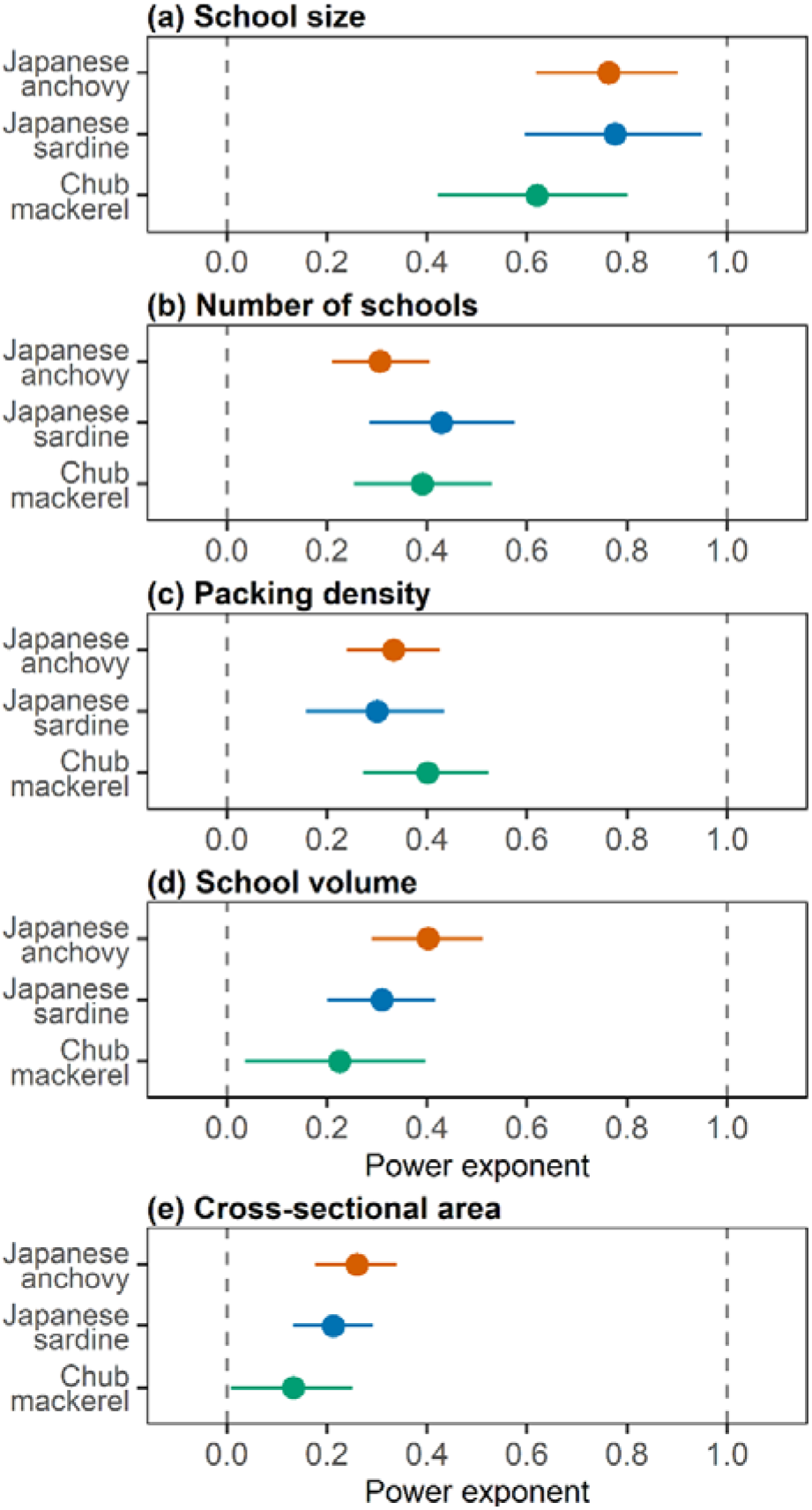
Estimated power exponents for the relationship between local population density and (a) school size, (b) the number of schools, (c) school packing density, (d) school volume, and (e) school cross-sectional area. Points indicate estimated values (posterior means), and error bars indicate 90% credible intervals.

**Fig. 3.**
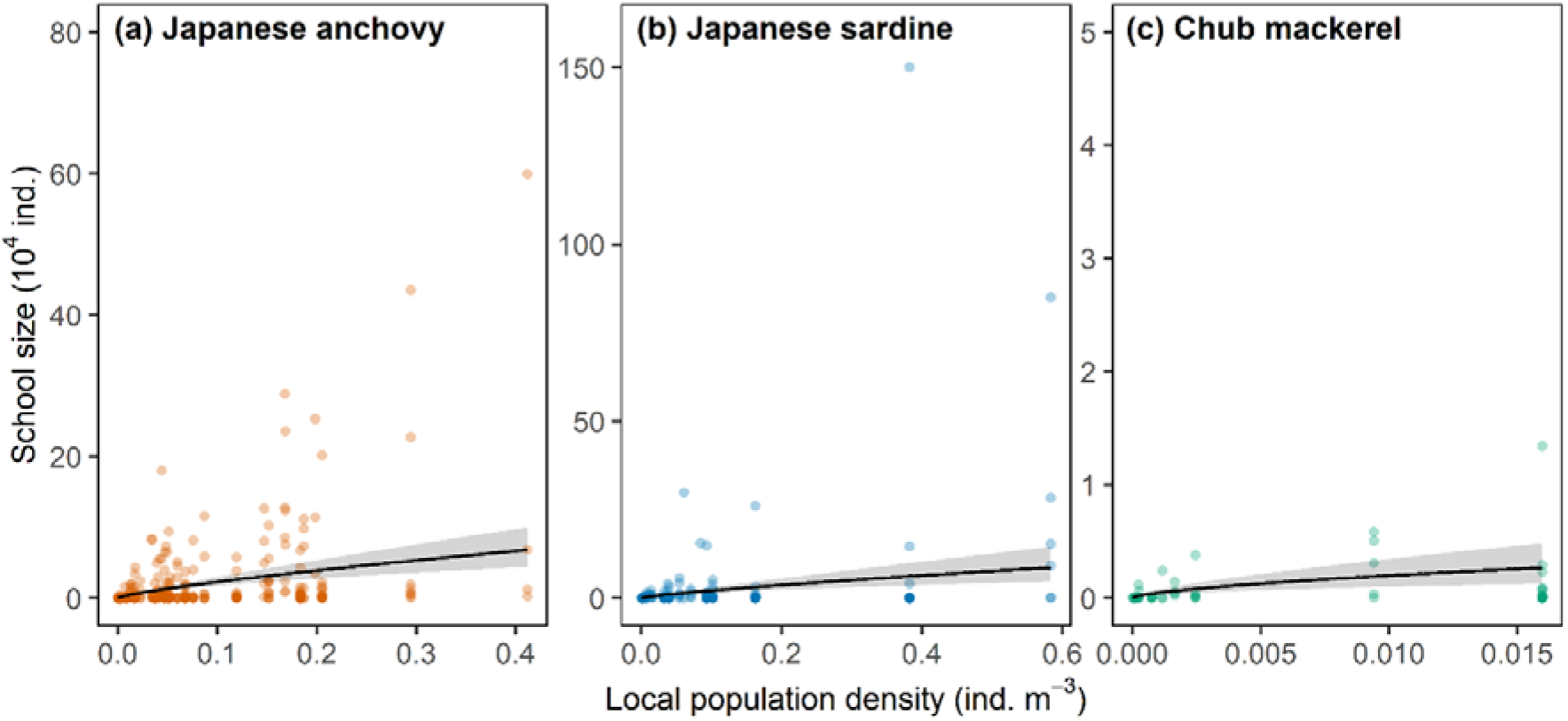
Relationship between local population density and school size for (a) Japanese anchovy, (b) Japanese sardine, and (c) chub mackerel. Translucent circles indicate observed values of individual schools. Solid lines indicate estimated relationships (posterior means), and grey bands indicate 90% credible intervals.

### Number of schools

For all species, the number of schools per km tended to increase as local population density increased, but the rate of increase slowed with increasing local population density (Fig. 4). For all species, the power exponents were greater than zero (indicating positive relationships) but much less than one (ranging from 0.31 to 0.43; Fig. 2b and Table S1), indicating that the rate of increase of the number of schools slowed considerably with increasing local population density (Fig. 4). The model with species-specific relationships between the number of schools and population density had a lower WBIC than the model with species-common relationships (species-specific, 222.4; species-common, 225.7), indicating that the relationships differed among species (Fig. S1b).

**Fig. 4.**
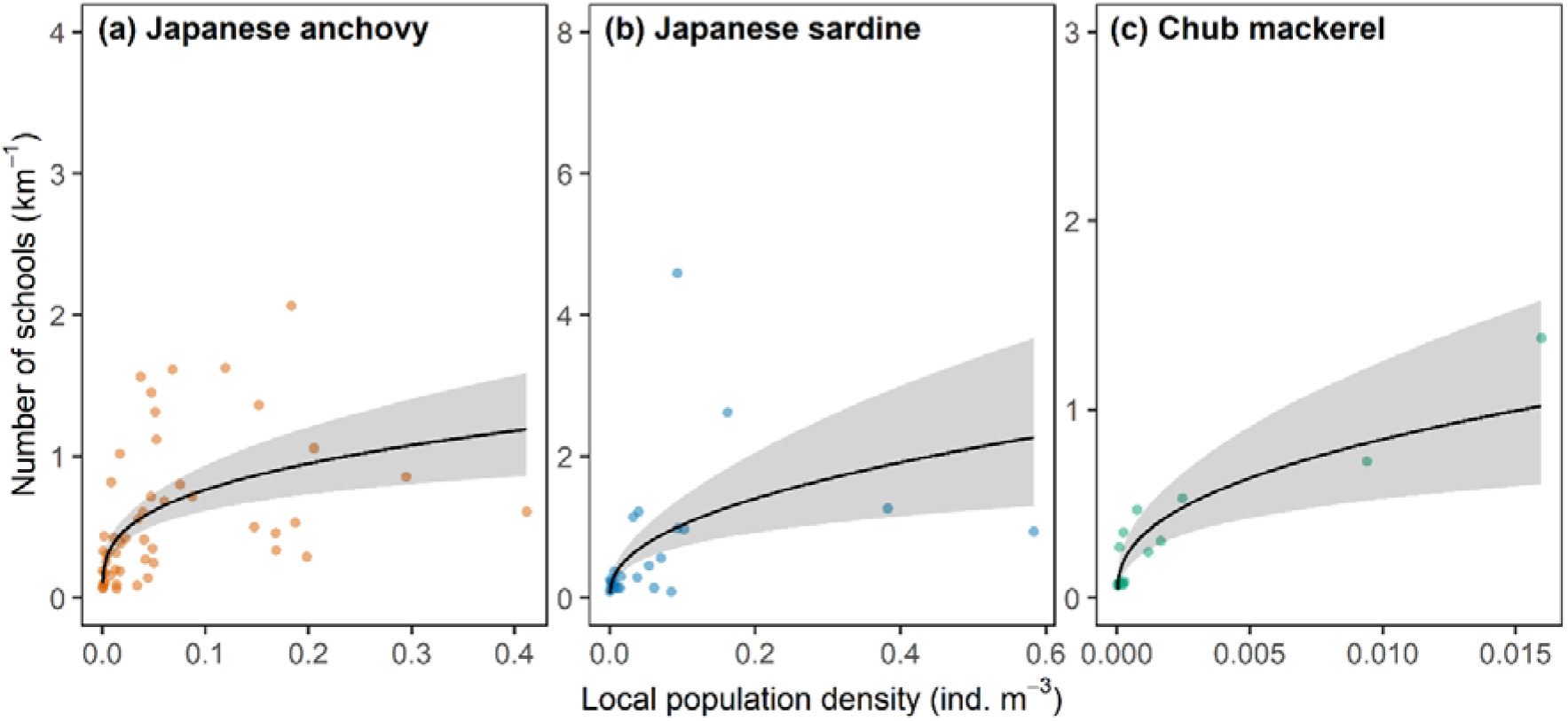
Relationship between local population density and the number of schools for (a) Japanese anchovy, (b) Japanese sardine, and (c) chub mackerel. Translucent circles indicate observed values at individual stations. Solid lines indicate estimated relationships (posterior means), and grey bands indicate 90% credible intervals.

### School packing density

In all species, packing density tended to increase as local population density increased, but the rate of increase slowed with increasing population density (Fig. 5). For all species, the power exponents were greater than zero (indicating positive relationships) but much less than one (ranging from 0.30 to 0.40; Fig. 2c and Table S1), indicating that the rate of increase of packing density slowed considerably with increasing local population (Fig. 5). The model with species-specific relationships between packing density and population density had a lower WBIC than the model with species-common relationships (species-specific, 456.1; species-common, 460.2), indicating that the relationships differed among species (Fig. S1c).

**Fig. 5.**
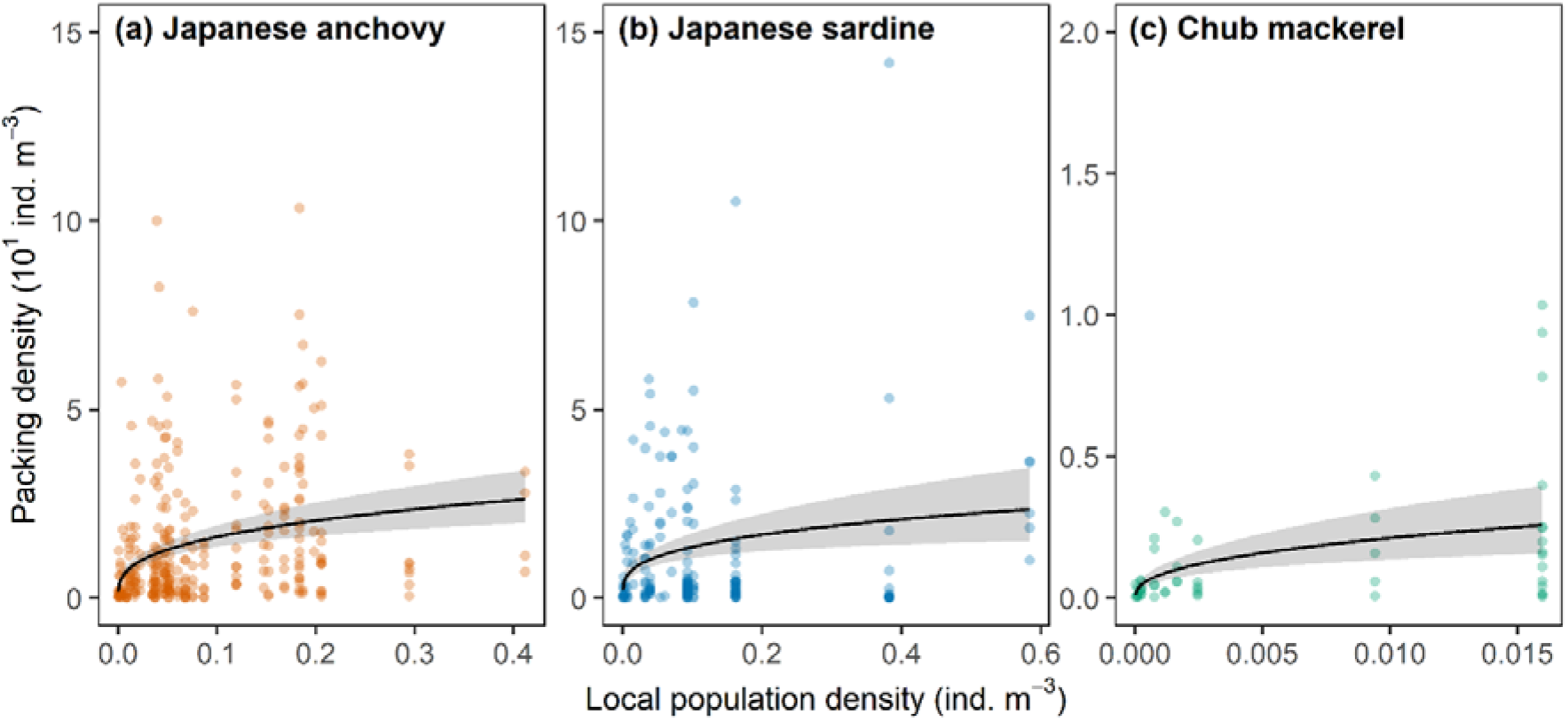
Relationship between local population density and school packing density for (a) Japanese anchovy, (b) Japanese sardine, and (c) chub mackerel. Translucent circles indicate observed values of individual schools. Solid lines indicate estimated relationships (posterior means), and grey bands indicate 90% credible intervals.

### Spatial extent of a school

For all species, school volume tended to increase with increasing local population density, but the rate of increase slowed as population density increased (Fig. 6). For all species, the power exponents were greater than zero (indicating positive relationships) but much less than one (ranging from 0.22 to 0.40; Fig. 2d and Table S1), indicating that the rate of increase of school volume slowed considerably with increasing local population (Fig. 6). The model with species-specific relationships between school volume and population density had a lower WBIC than the model with species-common relationships (species-specific, 208.1; species-common, 210.4), indicating that the relationships differed among species (Fig. S1d).

**Fig. 6.**
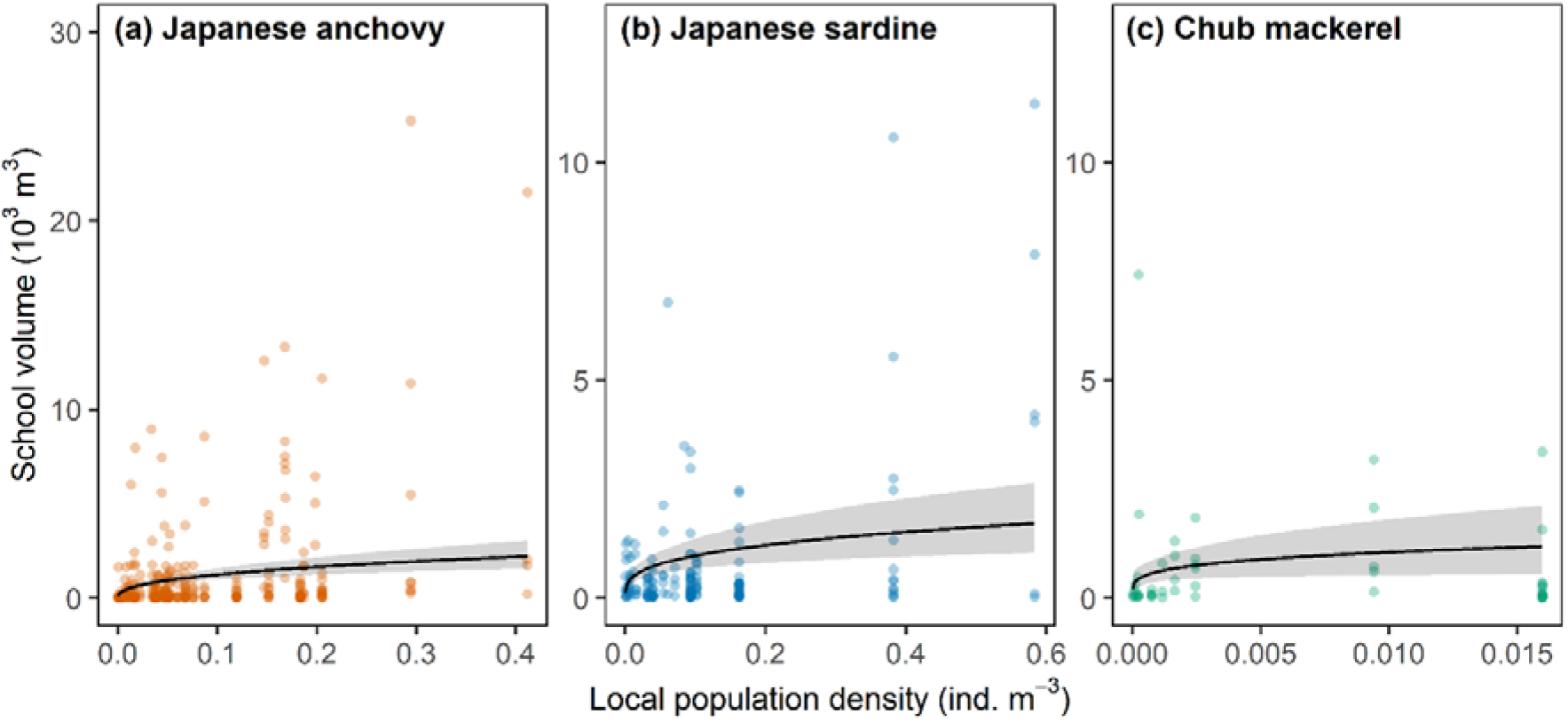
Relationship between local population density and school volume for (a) Japanese anchovy, (b) Japanese sardine, and (c) chub mackerel. Translucent circles indicate observed values of individual schools. Solid lines indicate estimated relationships (posterior means), and grey bands indicate 90% credible intervals.

In school cross-sectional area, for all species, the power exponents were greater than zero (indicating positive relationships) but much less than one (ranging from 0.13 to 0.26; Fig. 2e and Table S1), indicating that the rate of increase of cross-sectional area slowed considerably with increasing local population (Fig. 7). The model with species-specific relationships between cross-sectional area and population density had a lower WBIC than the model with species-common relationships (species-specific, 690.7; species-common, 693.1), indicating that the relationships differed among species (Fig. S1e).

**Fig. 7.**
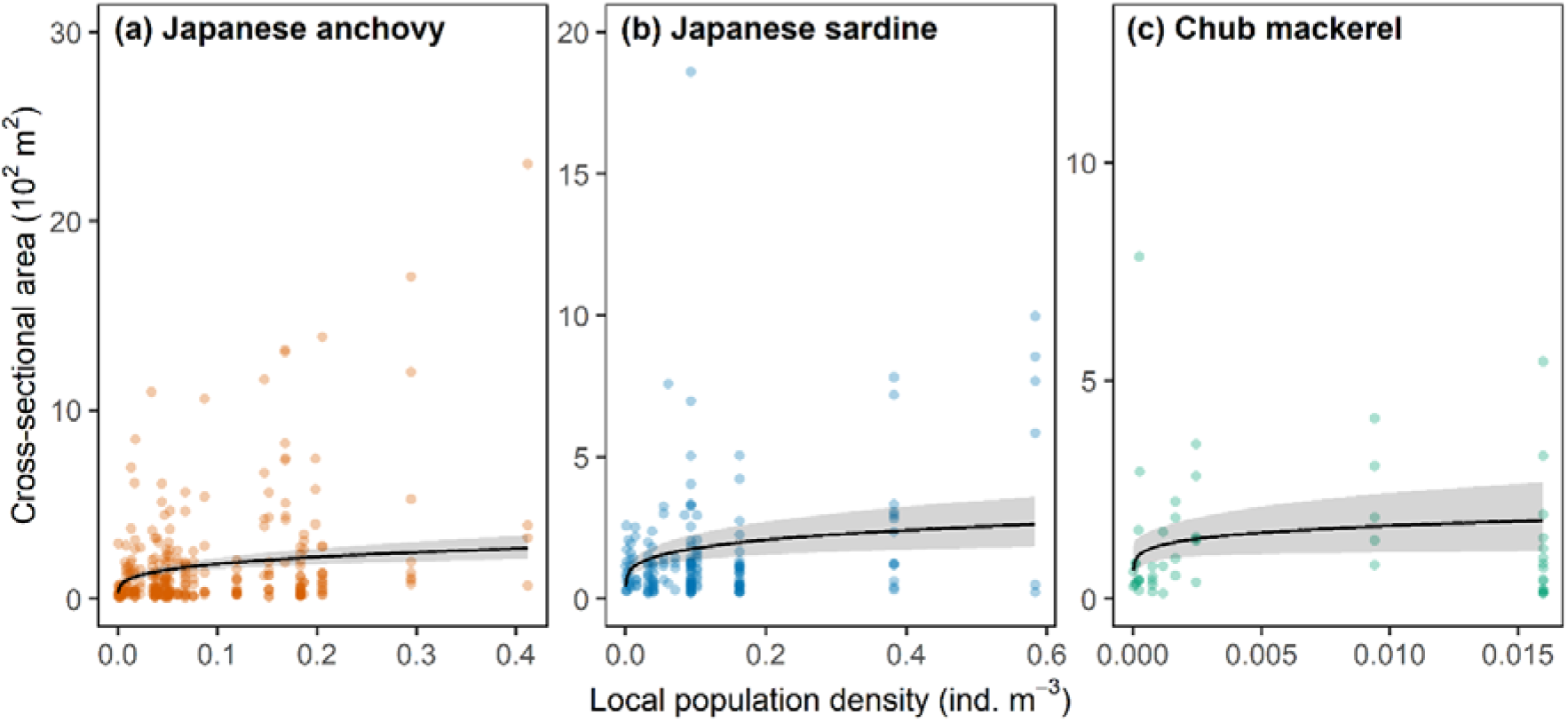
Relationship between local population density and school cross-sectional area for (a) Japanese anchovy, (b) Japanese sardine, and (c) chub mackerel. Translucent circles indicate observed values of individual schools. Solid lines indicate estimated relationships (posterior means), and grey bands indicate 90% credible intervals.

## Discussion

The effects of population density on school size and the number of schools is important for understanding population dynamics, community structure, and energy flow in marine ecosystems because the school is a fundamental ecological unit in which biological interactions such as competition and predation take place (Maury and Poggiale 2013; Maury 2017). However, information on the relationships between population density and school characteristics among small pelagic fishes in the field remains scarce, and quantitative relationships are yet to be clarified. In the present study, we show that school size increases almost linearly with population density, whereas the number of schools increases non-linearly in such a way that the rate of increase slows considerably with increasing population density. Our study provides important information on the quantitative relationships between population density and school characteristics.

Our results are largely consistent with the predictions of a theoretical model based on simple assumptions of encounter, merge, and split processes (Niwa 2004). In the model, schools are assumed to move at random through their environment. Schools merge whenever they encounter another school, and each school splits periodically into two schools. The schooling process of the studied species may resemble this simple model. However, there is a slight difference between our results and the predictions of the model. Whereas the model predicts that mean school size will increase linearly with population density, our results show that the rate of increase of mean school size slowed slightly with increasing local population size (i.e., a near-linear relationship), implying the existence of an optimal (stable) school size. The reason for this difference is likely that the theoretical model is purely mechanistic and does not consider the functions (i.e., the ultimate factors) of schooling behaviour. The formation of schools involves not only benefits but also costs, leading to the existence of an optimal group size that allows individuals to maximize their net benefit (Krause and Ruxton 2002; Ward and Webster 2016), and it has been suggested that individuals adjust their schooling behaviour to increase their net benefit (Sibly 1983). Accordingly, in our study species, the probability of school merging and splitting likely changed with school size in a way that served to increase this net benefit (Gueron and Levin 1995; Niwa 1998). A combination of mechanistic and functional approaches will be needed in future studies to better understand the dynamics of school size (Sumpter 2010; Griesser et al. 2011).

Our results show that whereas at low population densities, increasing density resulted in both more and larger schools, at high population densities, increasing density resulted in larger schools rather than more schools. This can be explained by the following mechanism: at low population densities, the frequency of encounters with conspecific schools is low, and hence many small schools are formed, whereas at high population densities, the frequency of encounter with conspecific schools is high, and as many schools merge to form larger schools, the number of schools becomes relatively stable. Similar trends have also been found in freshwater cyprinid fishes and terrestrial mammals, which form much smaller groups than marine small pelagic fishes (Southwell 1984; Pépin and Gerard 2008; Holubová et al. 2019). This suggests that this pattern might be widespread in animals that form fluid groups.

Our results also show that school packing density increased with increasing population density. This suggests that school size and packing density increase simultaneously as population densities increase. In fact, school size and packing density were strongly correlated (Japanese anchovy: ρ = 0.79, *p* < 0.001; Japanese sardine: *ρ* = 0.82, *p* < 0.001; chub mackerel: ρ = 0.75, *p* < 0.001). For small to mid-sized schools, predation risk per individual is expected to decrease as school size increases (Lima 1995; Krause and Ruxton 2002; Caro 2005). For large schools, however, as school size increases, the risk of predator attacks also increases as the school becomes more visible to predators (Pitcher and Parrish 1993; Maury 2017). Denser schools can engage in highly coordinated antipredator reactions through the efficient transmission of information within schools (Rieucau et al. 2014a, b), and therefore, the formation of dense schools could be a response of individuals against the threat of predation. Small pelagic fishes form especially massive schools, and individuals might cope with the increased risk of predator attacks at these large school sizes by forming dense and compact schools that allow them to efficiently obtain information from other schooling individuals.

In small pelagic fishes, a strong density-dependent decline in body growth is often observed during times of increasing population density (Wada et al. 1995; Watanabe and Yatsu 2004; Canales et al. 2020; Kamimura et al. 2021b). Density-dependent growth reduction is mainly caused by intraspecific competition for food (Hansen et al. 1999; Jenkins et al. 1999; Post et al. 1999; Kamimura et al. 2021b). Food competition can occur between schools and within a school, but within-school competition is likely to be stronger and more direct than between-school competition. Several studies in animals that form groups (although none of the studies were conducted on fish) have reported that competition for food within groups is more intense than competition between groups (Janson 1985, 1988). The increase in school size and packing density with increasing population density observed in our study is likely to create fairly strong competition for food. This might be one of the reasons for the strong negative density-dependence of growth observed in small pelagic fishes.

In our study, the shapes of the relationships between population density and school characteristics differed among small pelagic fishes (although the overall trends were similar across species). In krill, which are similar to small pelagic fishes in that they often form massive swarms and are preyed upon by a wide variety of pelagic megafauna, it has been reported that increasing population density causes a linear increase in the number of schools, but does not affect school size (Brierley and Cox 2015). This is different from the relationships observed in small pelagic fishes in our study. Differences in selection pressures (e.g., competition and predation) can lead to differences in optimal school size and hence to differences in aggregation tendency (i.e., the frequency of school fission and fusion) (Rieucau et al. 2015). This may account for the different shapes of the relationships between population density and school characteristics among species and taxa.

Furthermore, relationships between population density and school characteristics could also vary within a species depending on several factors. For example, seasonality (e.g., spawning and feeding seasons), body size, and age can affect optimal school sizes and aggregation tendencies (Hoare et al. 2000; Croft et al. 2003; Makris et al. 2009). Our study was conducted during the feeding season, and variations of body size and age in the study area were small throughout the study period (2007–2021). The body lengths of most chub mackerel and Japanese sardine corresponded to those of young of the year (Kamimura et al. 2021b, a). Further studies across seasons, locations, body sizes, ages, and species are needed to identify general patterns in relationships between population density and school characteristics.

Why small pelagic fishes form such massive schools remains an open question (Rieucau et al. 2015). In general, reducing predation risk is thought to be the most widely-applicable explanation for animal aggregation, although there are other explanations such as facilitating encounters with reproductive partners, improving foraging efficiency, and hydrodynamic gains (Pitcher and Parrish 1993; Krause and Ruxton 2002; Ward and Webster 2016). However, current theories on the function of schooling behaviour do little to explain the formation of massive schools (Rieucau et al. 2015; Maury 2017). This is because for massive schools, the benefits of schooling are likely to stay constant with increasing group size while the costs increase continually, meaning that the benefit-to-cost ratio declines rapidly with increasing group size and is quite small. The type of predator encountered has been suggested to be an important factor when assessing the benefits of massive schools (Rieucau et al. 2015), because the benefits of group size could be affected by predator hunting strategies (e.g., solitary vs. social predators) and attack efficiencies (e.g., the number of prey caught per strike) (Rieucau et al. 2015). In our results, the relationships between population density and school size differed among species, and the observed school size tended to be smaller for chub mackerel than for Japanese sardine and Japanese anchovy. The main predators of chub mackerel could differ from those of Japanese sardine and Japanese anchovy. A more detailed examination of these factors could help explain why small pelagic fishes form such massive schools.

Our results will be useful for population-status assessment of these species. Population density was positively correlated with school characteristics, indicating that school characteristics could be used as an indicator of population status. However, the relationships were highly nonlinear (except for that of school size). Therefore, this nonlinearity should be taken into account when using school characteristics as an indicator of population status (Erisman et al. 2011).

In the present study, we inferred schooling characteristics from echosounder data. Echosounder signals provide snapshot information in two dimensions: vertically through the water column and horizontally with successive acoustic signals along the vessel track (Simmonds and MacLennan 2008). This means that we were able to sample only a small fraction of each school we identified. In fact, several schools were estimated to be quite small. A previous study has reported that there are no significant differences between schooling characteristics obtained from echosounders and multibeam sonar, which can image an entire school in three dimensions (Brierley and Cox 2015). Therefore, the overall trend in our results is likely to be robust. However, the use of methods that can obtain three-dimensional and/or continuous-monitoring data will enable more accurate and detailed analysis (Makris et al. 2006; Paramo et al. 2007) in the future and could provide new biological and ecological insights.

In conclusion, our results show that school size increased almost linearly with increasing population density, whereas the number of schools and other school characteristics (school packing density, school volume, and school cross-sectional area) increased non-linearly such that the rate of increase slowed considerably with increasing population density. Furthermore, we found that the shapes of the relationships between population density and school characteristics differed among species, although the overall trend was similar. These results indicate that the schooling behaviour of small pelagic fish species is density-dependent, and responses to changes in density differ among species. Our results will help advance understanding of how biological interactions such as intra- and inter-specific competition and predator–prey interactions mediate the density-dependent processes that underlie the population dynamics and community structure of small pelagic fishes in marine ecosystems (Maury 2017).

## Acknowledgements

The dataset analysed in this study was made possible by enormous efforts undertaken by numerous researchers, as well as the captains, officers, and crews of the training vessel *Hokuho-Maru*. We thank Dr. K. Abe for providing information on methods for analysing echosounder data. We also thank English-speaking professional editors from ELSS, Inc., for English proofreading.

## Author contribution

Sho Furuichi conceived of the study and performed data analysis. All authors contributed to data collection. The first draft of the manuscript was written by Sho Furuichi and all authors commented on previous versions of the manuscript.

## Declarations

Conflict of interest

The authors declare no competing interests.

