## Supplementary material for "Density-dependent attributes of schooling in small pelagic fishes": Fig. S1, Table S1

**Table S1.** Estimates (posterior means) and 90% credible intervals (CI) of all parameters of all models analysed in this study.

| Response variable | Model structure | Parameter | Estimate | 90% CI |
| --- | --- | --- | --- | --- |
| School size | Species-specific | $a_{anchovy}$ | 13.62 | 7.78–21.73 |
| (10^4^ ind.) |  | $a_{sardine}$ | 13.30 | 6.46–23.77 |
|  |  | $a_{mackerel}$ | 4.38 | 0.83–11.65 |
|  |  | $b_{anchovy}$ | 0.76 | 0.62–0.90 |
|  |  | $b_{sardine}$ | 0.78 | 0.59–0.95 |
|  |  | $b_{mackerel}$ | 0.62 | 0.42–0.80 |
|  |  | $\alpha_{anchovy}$ | 0.33 | 0.30–0.37 |
|  |  | $\alpha_{sardine}$ | 0.29 | 0.25–0.33 |
|  |  | $\alpha_{mackerel}$ | 0.37 | 0.29–0.47 |
|  |  | $\sigma_{r}$ | 0.78 | 0.61–0.91 |
|  | Species-common | $a$ | 13.26 | 9.10–18.65 |
|  |  | $b$ | 0.78 | 0.70–0.86 |
|  |  | $\alpha_{anchovy}$ | 0.33 | 0.30–0.37 |
|  |  | $\alpha_{sardine}$ | 0.29 | 0.25–0.33 |
|  |  | $\alpha_{mackerel}$ | 0.36 | 0.29–0.47 |
|  |  | $\sigma_{r}$ | 0.78 | 0.62–0.90 |
| Number of schools | Species-specific | $a_{anchovy}$ | 1.58 | 1.05–2.25 |
|  |  | $a_{sardine}$ | 2.90 | 1.53–4.98 |
|  |  | $a_{mackerel}$ | 5.88 | 1.84–12.88 |
|  |  | $b_{anchovy}$ | 0.31 | 0.21–0.41 |
|  |  | $b_{sardine}$ | 0.43 | 0.28–0.58 |
|  |  | $b_{mackerel}$ | 0.39 | 0.25–0.53 |
|  |  | $\varphi_{anchovy}$ | 4.71 | 2.40–8.76 |
|  |  | $\varphi_{sardine}$ | 2.98 | 1.38–5.57 |
|  |  | $\varphi_{mackerel}$ | 8.60 | 2.62–21.94 |
|  |  | $\sigma_{r}$ | 0.23 | 0.02–0.48 |
|  | Species-common | $a$ | 1.64 | 1.22–2.14 |
|  |  | $b$ | 0.29 | 0.23–0.35 |
|  |  | $\varphi_{anchovy}$ | 4.12 | 2.26–7.12 |
|  |  | $\varphi_{sardine}$ | 2.76 | 1.35–4.87 |
|  |  | $\varphi_{mackerel}$ | 5.77 | 2.06–13.59 |
|  |  | $\sigma_{r}$ | 0.19 | 0.02–0.41 |

**Table S1.** (continued)

| Response variable | Model structure | Parameter | Estimate | 90% CI |
| --- | --- | --- | --- | --- |
| Packing density | Species-specific | $a_{anchovy}$ | 3.54 | 2.49–4.89 |
| (10^1^ ind. m^−3^) |  | $a_{sardine}$ | 2.79 | 1.67–4.32 |
|  |  | $a_{mackerel}$ | 1.51 | 0.53–3.21 |
|  |  | $b_{anchovy}$ | 0.33 | 0.24–0.43 |
|  |  | $b_{sardine}$ | 0.30 | 0.16–0.43 |
|  |  | $b_{mackerel}$ | 0.40 | 0.27–0.52 |
|  |  | $\alpha_{anchovy}$ | 0.84 | 0.74–0.94 |
|  |  | $\alpha_{sardine}$ | 0.60 | 0.51–0.70 |
|  |  | $\alpha_{mackerel}$ | 0.90 | 0.67–1.17 |
|  |  | $\sigma_{r}$ | 0.47 | 0.32–0.61 |
|  | Species-common | $a$ | 4.28 | 3.24–5.54 |
|  |  | $b$ | 0.44 | 0.37–0.50 |
|  |  | $\alpha_{anchovy}$ | 0.84 | 0.74–0.94 |
|  |  | $\alpha_{sardine}$ | 0.61 | 0.52–0.70 |
|  |  | $\alpha_{mackerel}$ | 0.75 | 0.56–0.99 |
|  |  | $\sigma_{r}$ | 0.59 | 0.46–0.72 |
| School volume | Species-specific | $a_{anchovy}$ | 3.19 | 2.07–4.71 |
| (10^3^ m^3^) |  | $a_{sardine}$ | 2.04 | 1.17–3.28 |
|  |  | $a_{mackerel}$ | 3.78 | 0.69–10.00 |
|  |  | $b_{anchovy}$ | 0.40 | 0.29–0.51 |
|  |  | $b_{sardine}$ | 0.31 | 0.20–0.42 |
|  |  | $b_{mackerel}$ | 0.22 | 0.04–0.40 |
|  |  | $\alpha_{anchovy}$ | 0.57 | 0.50–0.64 |
|  |  | $\alpha_{sardine}$ | 0.71 | 0.60–0.82 |
|  |  | $\alpha_{mackerel}$ | 0.64 | 0.49–0.82 |
|  |  | $\sigma_{r}$ | 0.83 | 0.74–0.92 |
|  | Species-common | $a$ | 2.13 | 1.55–2.85 |
|  |  | $b$ | 0.26 | 0.20–0.33 |
|  |  | $\alpha_{anchovy}$ | 0.57 | 0.50–0.64 |
|  |  | $\alpha_{sardine}$ | 0.70 | 0.60–0.82 |
|  |  | $\alpha_{mackerel}$ | 0.62 | 0.47–0.79 |
|  |  | $\sigma_{r}$ | 0.83 | 0.75–0.92 |

**Table S1.** (continued)

| Response variable | Model structure | Parameter | Estimate | 90% CI |
| --- | --- | --- | --- | --- |
| Cross-sectional area | Species-specific | $a_{anchovy}$ | 3.39 | 2.50–4.48 |
| (10^2^ m^2^) |  | $a_{sardine}$ | 2.96 | 2.00–4.18 |
|  |  | $a_{mackerel}$ | 3.45 | 1.20–7.06 |
|  |  | $b_{anchovy}$ | 0.26 | 0.18–0.34 |
|  |  | $b_{sardine}$ | 0.21 | 0.13–0.29 |
|  |  | $b_{mackerel}$ | 0.13 | 0.01–0.25 |
|  |  | $\alpha_{anchovy}$ | 1.09 | 0.95–1.23 |
|  |  | $\alpha_{sardine}$ | 1.31 | 1.11–1.54 |
|  |  | $\alpha_{mackerel}$ | 1.31 | 0.99–1.70 |
|  |  | $\sigma_{r}$ | 0.57 | 0.46–0.68 |
|  | Species-common | $a$ | 2.71 | 2.19–3.32 |
|  |  | $b$ | 0.17 | 0.13–0.22 |
|  |  | $\alpha_{anchovy}$ | 1.08 | 0.95–1.22 |
|  |  | $\alpha_{sardine}$ | 1.30 | 1.11–1.53 |
|  |  | $\alpha_{mackerel}$ | 1.26 | 0.96–1.64 |
|  |  | $\sigma_{r}$ | 0.57 | 0.46–0.68 |

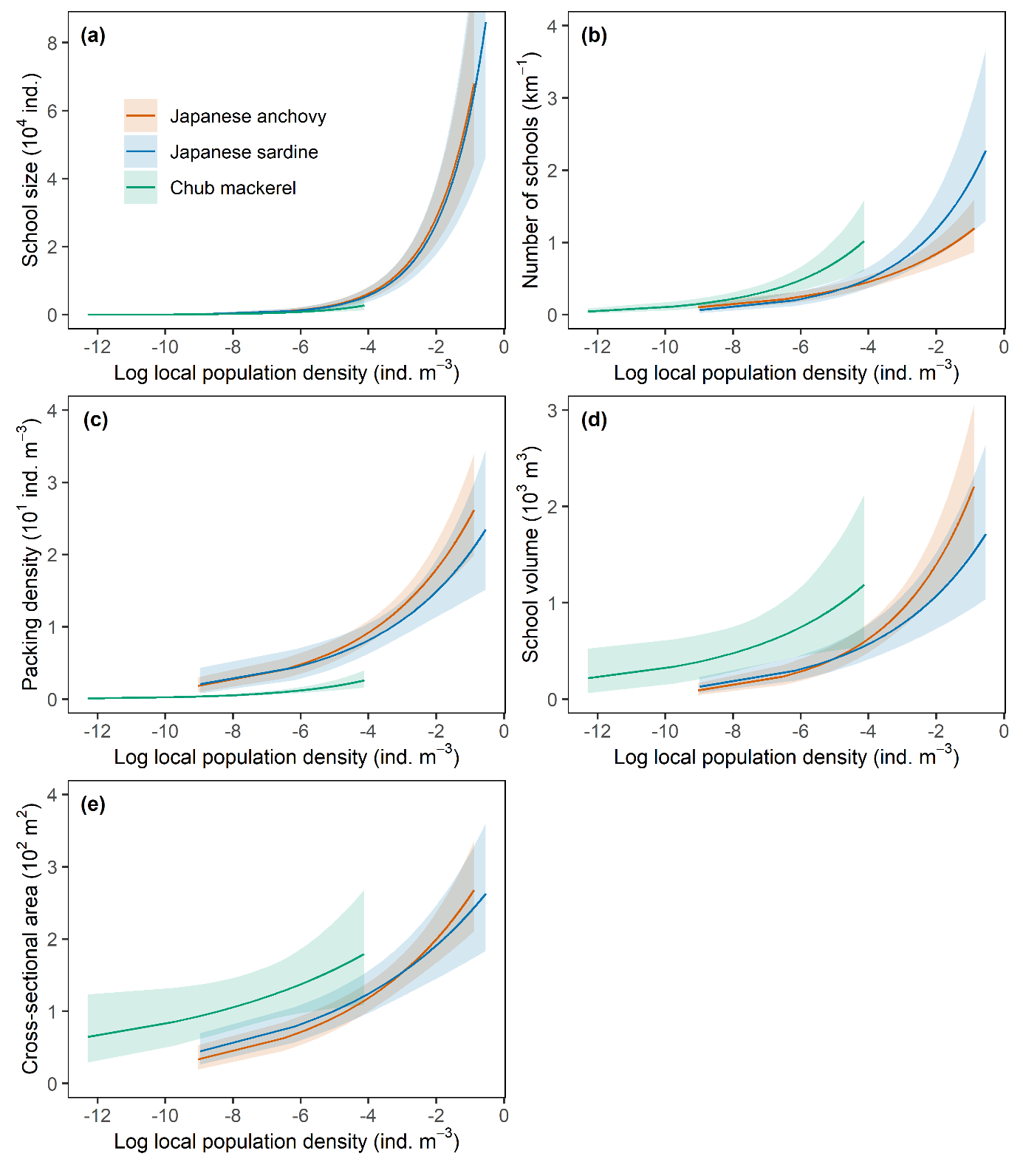

**Fig. S1**	Comparison of relationships between local population density and (a) school size, (b) the number of schools, (c) school packing density, (d) school volume, and (e) school cross-sectional area among fish species. Note that the horizontal axes are log-transformed to allow comparison among species with varied ranges of local population density. Solid lines indicate estimated relationships (posterior means), and shaded bands indicate 90% credible intervals.
